# Species interactions determine plasmid persistence in a 3-member bacterial community

**DOI:** 10.1101/2025.05.31.657151

**Authors:** Kaitlin A. Schaal, Ying-Jie Wang, Johannes Nauta, Manlio De Domenico, Shai Pilosof, James P. J. Hall

**Author notes:** **Author contributions** Conceptualization: all authors. Data curation: KS. Formal analysis: KS, JPJH. Funding acquisition: MdeD, SP, JPJH. Investigation: KS. Methodology: KS. Project administration: MdeD, SP, JPJH. Supervision: MdeD, SP, JPJH. Visualization: KS. Writing — original draft: all authors. Writing — review & editing: all authors.

## Abstract

Microbial communities are shaped by multiple forces, including interspecies interactions and the effects of mobile genetic elements such as plasmids. How these forces interact to affect community responses to environmental perturbations remains unclear, particularly considering the qualitatively different natures and the complexity of competitive and plasmid transfer interactions. We investigated the role of bacterial and plasmid interaction networks in community responses to single and multiple environmental perturbations, using a model community of three bacteria and two plasmids grown in rich media over five 2-day transfers. Bacterial interactions were the primary driver of community response, to the extent that plasmids were not always retained in the community even when they carried relevant resistance genes. Overall, our results indicate that while bacterial and plasmid interactions both shape community responses to environmental perturbations, bacteria-bacteria interactions may be the primary driver of community dynamics and their response to perturbations.

## Introduction

Microbial communities are governed by diverse interactions. Microbes can interact antagonistically, competing for the limited resources necessary for growth and replication, or producing factors that kill or impede the growth of competitors (1). They can also interact mutualistically, facilitating one another’s growth by cross-feeding and cooperation to use resources (2–5). However, interactions in microbiomes are not restricted to interactions between cellular organisms. Superimposed on the various species- and strain-level interactions are interactions between cells and their cognate mobile genetic elements (MGEs), which include bacteriophages and plasmids (6). Through such interactions, MGEs can exert significant effects on a community. MGEs can act as parasites, burdening community members in a manner that can prevent a species or strain from dominating (e.g., Kill-The-Winner dynamics in bacteriophages) (7,8). Conversely, MGEs can confer context-dependent beneficial traits on the microbes that host them, including resistance to environmental perturbations like antibiotics or heavy metals, the ability to metabolize novel resources, or traits allowing colonization of new niches (9,10).

Central to MGE-microbe interactions is the concept of host range: the ability of an MGE to enter and replicate within a given recipient cell type (its ‘host’). In the context of beneficial MGE-host interactions, strains that are susceptible to acquiring MGEs can rapidly gain new traits which might accelerate adaptation to changing environments. Hosts of MGEs act as reservoirs of adaptive traits that can be disseminated to others in the community (11), enhancing community-wide adaptation, and strains which can acquire multiple MGEs (‘bridge’ strains) and thereby enable recombination between MGEs could enable new combinations of functionality (12,13). However, bridge strains may suffer excessively from metabolic and other burdens associated with carriage of multiple MGEs, ultimately resulting in their out-competition by other organisms and local extinction (14,15), with potential consequences for MGE persistence. Bridge strains are therefore expected to face a trade-off shaped by the costs of multiple plasmids versus the environment-dependent benefits of plasmids’ genetic cargo.

To experimentally address how the presence of a bridge strain affects community dynamics and the persistence of MGEs, we developed a simple microbial community consisting of three bacterial strains (*Pseudomonas fluorescens* SBW25, *Escherichia coli* MG1655, and *P. putida* KT2440) and two plasmids (pQBR57, which confers mercury resistance, and pKJK5, which confers kanamycin resistance) (Figure 1A). Under our experimental conditions, each plasmid can be hosted by two strains: pQBR57 by *P. fluorescens* SBW25 and *P. putida* KT2440, and pKJK5 by *E. coli* MG1655 and *P. putida* KT2440. As *P. putida* KT2440 can host both plasmids (and can harbour both at the same time) we consider it a ‘bridge’ strain, and the other two as ‘peripheral’ strains. We tracked bacterial and plasmid populations, under non-selective conditions, selective levels of mercury, selective levels of kanamycin, and under combined selection from both, for ∼38 generations. We compared the results with communities lacking plasmids, lacking the bridge strain, or lacking both, to ascertain the contribution of each member to community dynamics (Figure 1B). We found that the presence of plasmids increased community resilience in the face of environmental stresses. However, bacterial species interactions played a predominant role in the dynamics, occasionally driving plasmid extinction even under positive selection.

**Figure 1.**
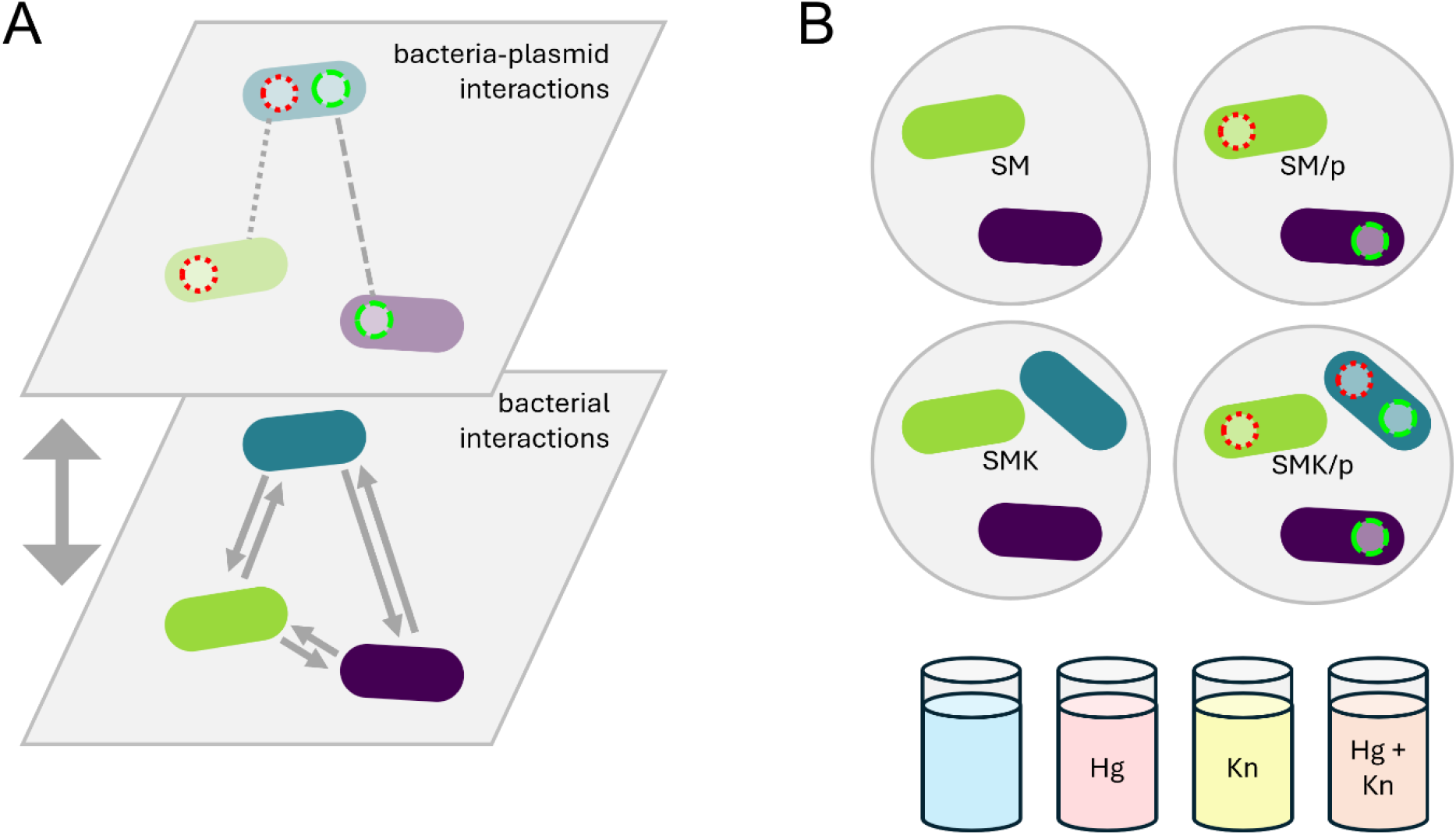
Experimental design. The community interaction network (A) consists of three bacterial strains (lower layer) and two plasmids (upper layer). The bacteria are *P. fluorescens* SBW25 (green oval), *P. putida* KT2440 (blue oval), and *E. coli* MG1655 (purple oval). The plasmids are pQBR57 (red dotted ring) and pKJK5 (green dashed ring). KT2440 can host both plasmids, making it a bridge strain within the community. We analysed dynamics of four communities under four environmental conditions, including three stress treatments (B).

## Methods

### Bacterial strains and plasmids

We used *Pseudomonas fluorescens* SBW25, *Pseudomonas putida* KT2440, and *Escherichia coli* MG1655. SBW25 (16,17) and the plasmid pQBR57 (18–20) were isolated independently from a sugar beet field in Oxfordshire, UK. KT2440 is a derivative of *P. putida* mt-2 (21). MG1655 is a derivative of *E. coli* K-12 (22). Interactions between SBW25 and KT2440 have been previously studied, as well as dynamics between the two strains involving pQBR57, which carries the *merA* gene for mercury resistance (11,23). pQBR57 is considered a narrow host range plasmid, as transconjugants have only been isolated using Pseudomonads (23). pKJK5 is a broad host range plasmid known to infect both KT2440 and various *E. coli* strains (24,25) including MG1655 (26). Though pKJK5 has been reported to conjugate into SBW25 (27), we have been unable to retrieve SBW25 (pKJK5) transconjugants either under the experimental conditions outlined in this study, or an extended range of conditions including broth and filter mating, and selection for pKJK5-encoded traits. We therefore consider SBW25 as incapable of hosting pKJK5 in the context of our experimental system. Both plasmids are relevant for soil microbial communities (23,26). Strains were cultured in LB (10 g/L peptone from casein, 5 g/L yeast extract, 10 g/L NaCl, with agar at 1.5% where required) and KB (King’s B; 20 g/L proteose peptone, 1.5 g/L MgSO_4_, 1.15 g/L K_2_HPO_4_, 10 g/L glycerol, with agar at 1.2% as required) as described.

We used derivatives of each strain to enable enumeration of different populations: SBW25-Sm^R^*lacZ* (23), KT2440-Gm^R^ (11), and wild-type MG1655 (which carries the *lacZ* gene). We used pQBR57::tdTomato (28) and pKJK5::GFP, which has the GFP gene inserted into the tetracycline resistance gene and is resistant to kanamycin but not tetracycline (26). This allowed us to track the three bacterial strains and plasmid carriage throughout the experiment: the plasmids by transfluorescence on a blue light box, and the strains by plating on agar with 250 µg/ml streptomycin and incubating at 28°C (for SBW25) and on agar with 50 µg/ml X-Gal and incubating at 37°C (for KT2440 and MG1655).

### Sub-population construction

For brevity, we named our different community treatments SMK (for SBW25 + MG1655 + KT2440) or SM (for SBW25 + MG1655) for communities containing or lacking the bridge strain respectively, appended with ‘/p’ for communities in which plasmids were initially present. Our experimental design (Figure 1) used a ‘link-balanced’ initial community composition (Table 1), whereby all populations in treatments including plasmids are initiated with 50% plasmid-free cells, with the remaining 50% evenly comprised of plasmid-carrying sub-populations (one each for SBW25 and MG1655, three for KT2440). We therefore constructed all possible donor strains (Table S1) prior to starting the experiment, using the patch conjugation method (29). To test effects of plasmid carriage on bacterial growth rate, we grew overnight cultures of each plasmid-free and plasmid-carrying strain, transferred them by pin replicator (∼1.5 µl volume) into 200 µl LB broth in a 96-well plate, and performed a 48-hr growth curve at 28 °C shaking in a Tecan Infinite F Nano+ plate reader (Tecan Group Ltd., Männedorf, Switzerland), reading OD_600_ every 15 minutes. To test the effect of environmental stressors, growth curves were also established under a range of mercury conditions (0, 2.5, 5, 10, 20 µM).

**Table 1.** Initial community compositions. Fraction refers to the initial proportion of cells from each population.

| Community | Population | Fraction | Sub-population | Fraction |
| --- | --- | --- | --- | --- |
| SM | SBW25 | 0.5 |  |  |
|  | MG1655 | 0.5 |  |  |
| SMK | SBW25 | 0.33 |  |  |
|  | MG1655 | 0.33 |  |  |
|  | KT2440 | 0.33 |  |  |
| SM/p | SBW25 | 0.5 | SBW25 | 0.25 |
|  |  |  | SBW25(pQBR57) | 0.25 |
|  | MG1655 | 0.5 | MG1655 | 0.25 |
|  |  |  | MG1655(pKJK5) | 0.25 |
| SMK/p | SBW25 | 0.33 | SBW25 | 0.165 |
|  |  |  | SBW25(pQBR57) | 0.165 |
|  | MG1655 | 0.33 | MG1655 | 0.165 |
|  |  |  | MG1655(pKJK5) | 0.165 |
|  | KT2440 | 0.33 | KT2440 | 0.165 |
|  |  |  | KT2440(pQBR57) | 0.055 |
|  |  |  | KT2440(pKJK5) | 0.055 |
|  |  |  | KT2440(pKJK5, pQBR57) | 0.055 |

### Experiment

We streaked out all strains on LB agar plates and incubated overnight at 28°C. We selected 6 independent colonies of each and grew them overnight in LB liquid at 28°C and 180 RPM in 50-ml Falcon tubes in an Innova 42 incubator shaker. We standardized overnight cultures to OD_600_ 1 in LB then diluted 10x, and assembled communities by combining strains in the ratios shown in Table S1 in a total volume of 600 µl. We prepared a 96-well plate with 180 µl LB per well and inoculated each replicate community four times (once per environmental treatment), to a total of 200 µl. Communities were started with 5 × 10^7^ CFU. The plating pattern was randomized by sorting the list of samples by the rand() function in Microsoft Excel. No environmental stresses were added during the first growth cycle. We incubated the experimental plate for 2 days at 28°C and 640 RPM on a Stuart Microtiter SSL-5 orbital shaker then transferred to the next cycle by Boekel Scientific pin replicator (transfer volume ∼1 µl). All subsequent 96-well plates were prepared with 200 µl LB per well, with ¼ of wells containing 10 µM HgCl2 [Hg(II)], ¼ containing 12.5 µg/ml kanamycin sulfate, and ¼ containing 10 µM HgCl2 [Hg(II)] and 12.5 µg/ml kanamycin sulfate; the remaining ¼ received no perturbation. Concentrations of mercury and kanamycin were the lowest relevant stress levels for the non-plasmid-hosting strains. We passaged the cultures 4 times over a total of 10 days. We quantified population and sub-populations of the initial communities and after cycles 1, 3, and 5 (days 2, 6, and 10), with a lower limit of detection of 5 × 10^5^, or >0.5% of community carrying capacity. With a transfer volume of 1% of the community, we expected sub-populations below our limit of detection to be ultimately removed from the community owing to dilution during transfers.

### Statistical analysis

We summarized overall dynamics of bacteria and plasmids for statistical analysis by calculating trapezoidal area under the curve (AUC, Figures 2 & 3), to avoid analytical complications that would have been introduced in time-series analysis by autocorrelation between timepoints (30–32). To quantify the effects of community structure and environmental stresses on bacterial and plasmid dynamics, we used general linear models followed by Tukey HSD tests. To quantify the extinction probability of SBW25, we ranked extinction as a binary event and performed a Fisher’s exact test. We analysed the data using R version 4.5.2 (33) and RStudio version 2026.01.1+403 (34). We organized the data using the “tidyr” (35) and “dplyr” (36) packages, analysed growth curve data with “gcplyr” (37), and visualized them using “ggplot2” (38) and “viridis” (39). All data and analysis files can be found at [public DOI after review].

**Figure 2.**
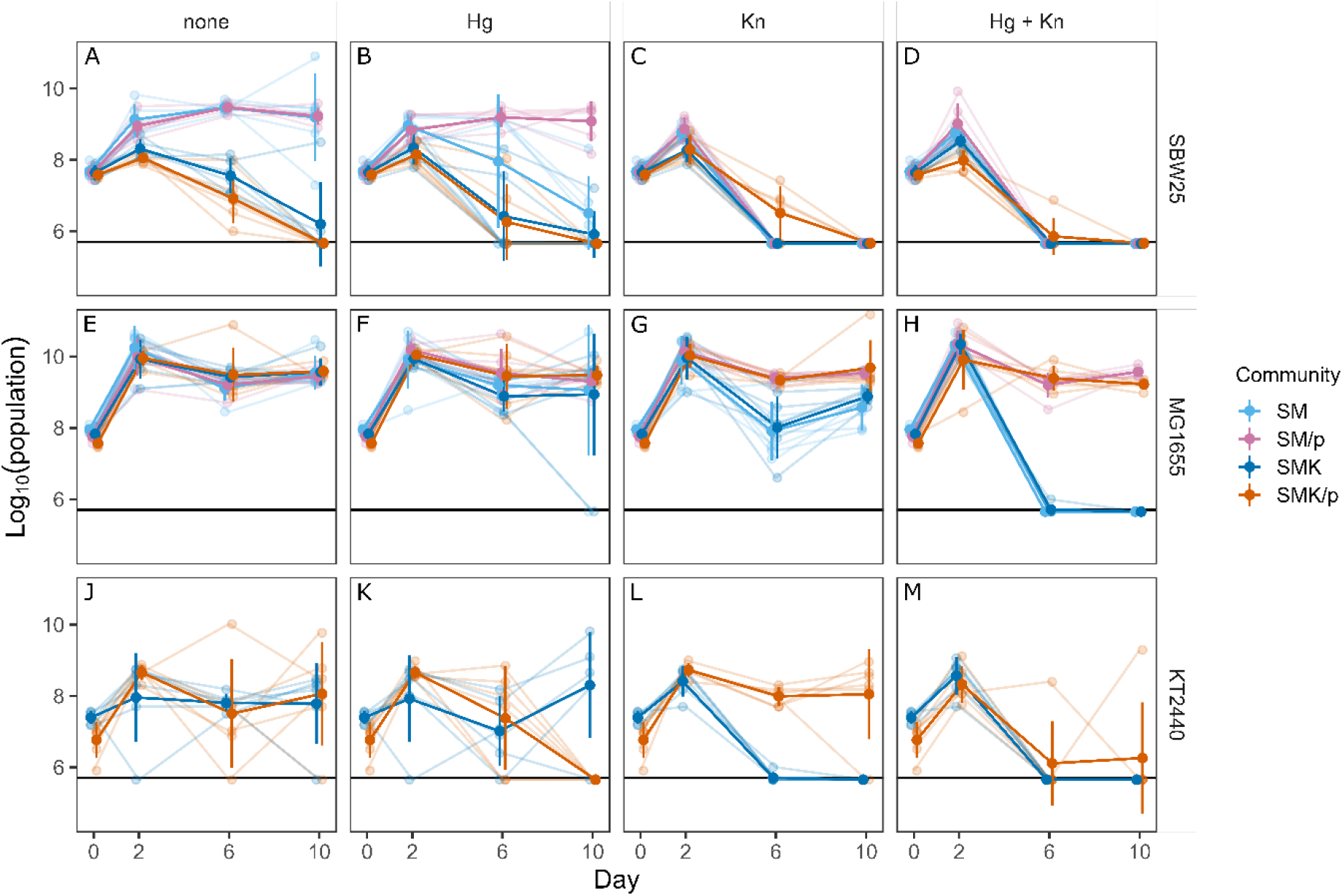
Bacterial dynamics. Population sizes of SBW25, KT2440, and MG1655 over 10 days under different environmental conditions (environmental perturbations were included from day 2 onward). The upper row of panels shows dynamics when plasmids were absent from the community, and the lower row shows bacterial dynamics when plasmids were present. Small dots represent replicate values, large dots represent means of 6 biological replicates, and vertical lines represent 95% confidence intervals of the means. Stresses were added after day 2.

**Figure 3.**
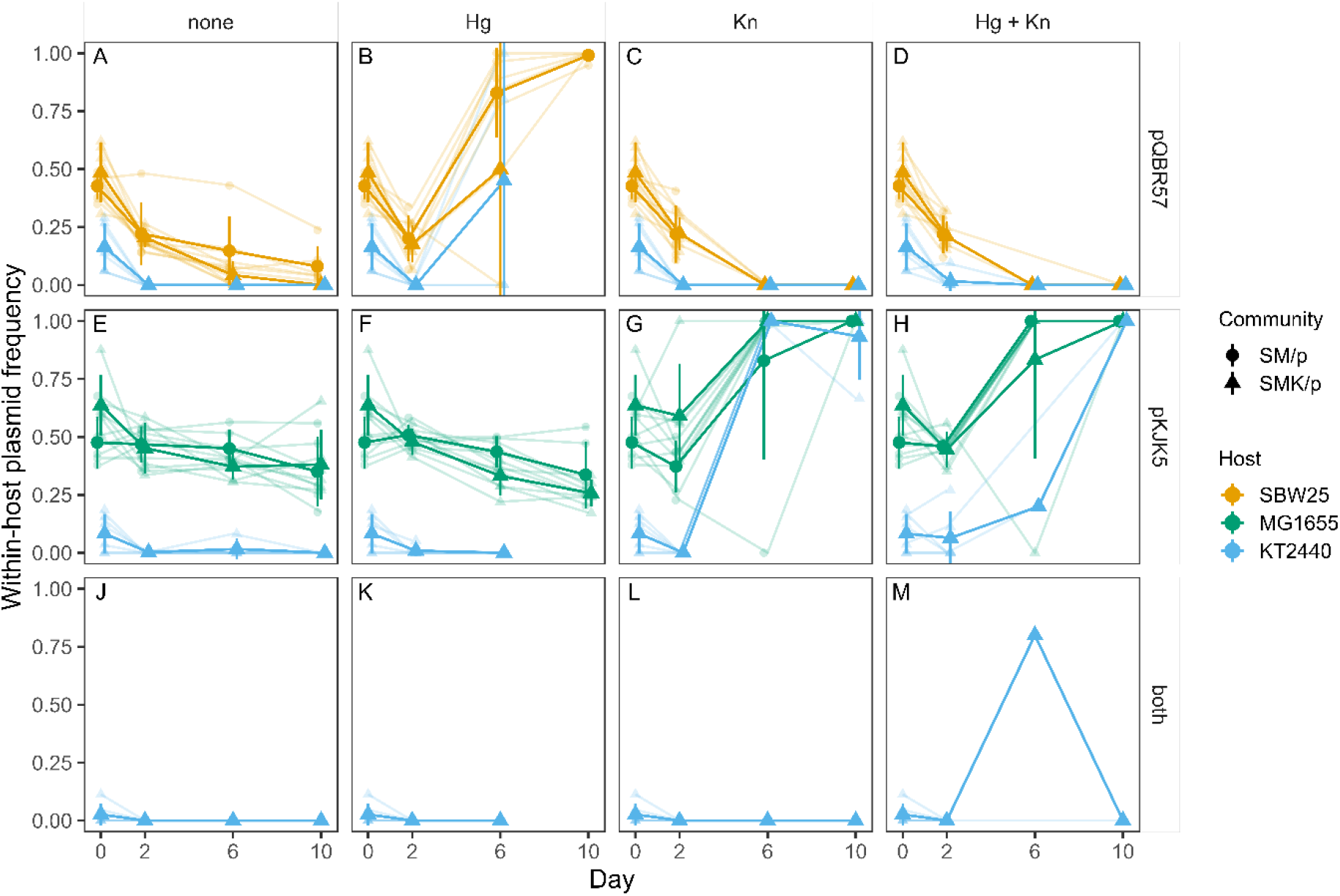
Plasmid frequency dynamics. Frequencies of plasmid-carrying subpopulations of the host bacteria over 10 days under different environmental conditions. pQBR57 confers Hg resistance and pKJK5 confers Kn resistance. A frequency of 1 indicates that all cells of that host contain the plasmid. Colours represent plasmid status (hosting one plasmid, the other, or both plasmids) and shapes represent host species. The upper row of panels shows plasmid frequency dynamics in the 2-member community, and the lower row shows dynamics in the 3-member community. Small dots represent replicate values, large dots represent means of 6 biological replicates, and vertical lines represent 95% confidence intervals of the means. Missing data indicates where hosts were no longer detected in the community, to differentiate from plasmid frequency of 0 within the host.

## Results

### Plasmids rescue total community population size from stress

To investigate the effect that plasmid presence has on the overall community, we first considered communities lacking the bridge strain. In the absence of plasmids, exogenous stresses (mercury, kanamycin, or both) decreased cumulative community size (Figure S1; ANOVA main effect of environment on AUC, F_[3,80]_ = 72.54, *p*-value < 0.0001), an effect principally driven by the impact of kanamycin (post-hoc Tukey tests effect of kanamycin *p*-values < 0.0002; effect of mercury *p*-value = 0.055). As expected, adding resistance-encoding plasmids to the communities ameliorated these effects (ANOVA main effect of plasmids, F_[1,80]_ = 173.4, *p*-value < 0.0001), effectively restoring pre-stress community population size (post-hoc Tukey tests, *p*-values > 0.89). Plasmid addition had no significant effect on total community size in the absence of stress (post-hoc Tukey test, *p*-value = 1). At the community level, therefore, plasmids provided robustness to stress.

### Plasmids stabilized the two-member community

Under non-stressful conditions, plasmid addition did not significantly alter the dynamics of either of the two community members (Figure 2A&E; AUC post-hoc Tukey HSD tests, *p*-value > 0.9 for all comparisons), possibly because the presence of plasmid-free subpopulations in each strain masked the negative effect of plasmid burden on the overall population counts. Stresses particularly impacted SBW25, which performed worse under all stress treatments (Figure 2A-D; post-hoc Tukey HSD tests, Hg *p* = 0.03, Kn and Hg + Kn *p*-values < 0.001). As predicted given the host ranges of the plasmids, for SBW25 plasmid addition mitigated only the effect of mercury stress (Figure 2B; SM community vs SM/p community, *p*-value = 0.0006), and could not rescue the population when faced with kanamycin stress (Figure 2C; SM community vs SM/p community, *p*-values = 1), where SBW25’s population fell below the detection limit of the experiment in each case.

In contrast, MG1655 was only significantly affected by kanamycin stress (Figure 2E-H, Hg *p* = 0.25, Kn and Hg + Kn *p*-values < 0.003), and though kanamycin significantly reduced MG1655’s population size (Figure 2G&H; post-hoc Tukey tests, *p*-value < 0.0001), the strain remained at appreciable levels even when plasmids were absent from the community (Figure 2G&H; AUC difference from non-stressful conditions of -7.34 for Kn vs. - 21.41 for Hg + Kn), possibly by emergence of a *de novo* mutant with increased kanamycin resistance. MG1655 clones isolated from plasmid-free populations show higher resistance to increased kanamycin concentrations than the ancestor, but lower than what is conferred by pKJK5 (Figure S2). Plasmid addition improved MG1655’s growth under kanamycin conditions, even rescuing MG1655 from dual stress (*p*-values < 0.0001).

In a two-member community with no between-species HGT, plasmids were therefore able to rescue their hosts from relevant stresses. Despite its inability to host the plasmid conferring mercury resistance, MG1655 demonstrated an ability to survive under mercury conditions regardless of plasmid presence, or under dual stress when it carries the plasmid conferring kanamycin resistance (Figure 2F&H). This may be due to mercury absorption by the growth medium and a higher overall tolerance to mercury than SBW25 and KT2440 (Supplemental results).

### Adding the bridge strain destabilized the community

We predicted that a bridge strain (Figure 1) will experience higher plasmid burden, altering its dynamics compared to when grown with no plasmids. We therefore investigated the dynamics of communities including a third member, *P. putida* KT2440, which can host both pQBR57 and pKJK5 and thus acts as a bridge strain when the plasmids are present.

First, we considered the effect of the bridge strain on plasmid-free communities. The presence of the bridge strain did not significantly influence total population size (Figure S1; ANOVA, F_[1,80]_ = 2.2, *p*-value = 0.13), and MG1655 was unaffected by KT2440’s presence across environmental treatments (Figure 2E-H; ANOVA, F_[1,80]_ = 0.095, *p*-value = 0.75). However, SBW25 was significantly impacted, experiencing population decline over the course of the experiment when the bridge strain was present (Figure 2A&B; ANOVA, main effect of bridge strain, F_[1,80]_ = 150.7 *p*-value < 0.0001). This difference in dynamics between the bacterial strains may have occurred due to niche overlap between the two *Pseudomonas* strains, resulting in nutrient competition (40). The effect of KT2440 on SBW25 was observed both in non-stressful and mercury conditions (Figure 2A&B; post-hoc Tukey HSD tests, *p*-values < 0.002) and drove SBW25 below the detection limit under non-stressful conditions (Figure 2A; Fisher’s Exact test, *p*-value = 0.004). We did not detect an effect in treatments containing kanamycin (*p*-values > 0.99) because SBW25 quickly fell below the detection limit regardless of KT2440 presence (Figure 2C&D). The presence of the bridge strain therefore shaped the overall community principally by influencing the dynamics and persistence of specific community members, in this case SBW25.

### Stress type determines plasmid effects on community stability

Given that plasmids stabilized the 2-member community and the bridge strain destabilized the plasmid-free community, we investigated how the presence of plasmids and bridge strain interacted to affect community members. Hosting pKJK5 imposes a cost on the bridge strain relative to hosting pQBR57 (Figure S3; ANOVA for differences in growth dynamics of KT2440 sub-populations, F_[3,23]_ = 3.86, *p*-value = 0.023; post-hoc Tukey HSD test for pKJK5 vs pQBR57, *p*-value = 0.034). However, we expected plasmids to facilitate growth of their hosts under conditions that selected for the cognate resistance traits.

Under mercury conditions, in contrast to the two-species communities, the presence of plasmids in the three-species communities did not improve survival of the pQBR57-hosting strains, SBW25 and KT2440. In fact, while there was no significant effect on SBW25 survival (Figure 2B; post-hoc Tukey HSD tests for SMK vs SMK/p, *p*-value >0.99), plasmid presence exerted a negative effect on KT2440, whose loss from the community was more likely when plasmids were present than when they were absent (Figure S4; Fisher’s Exact test, *p*-value = 0.015). This outcome suggests that the interaction between plasmids and environmental conditions can influence strain interactions in non-intuitive ways.

By contrast, pKJK5 hosts MG1655 and KT2440 grew better under kanamycin stress when plasmids were present. Plasmids improved MG1655 survival under both conditions containing kanamycin (Figure 2G&H; post-hoc Tukey HSD tests, *p*-value < 0.0001), while plasmids rescued KT2440 under kanamycin stress (Figure 2L; post-hoc Tukey HSD test, *p* < 0.001), but not dual stress, even though KT2440 is capable of hosting both plasmids and thus expressing resistances to both stresses (Figure 2M; post-hoc Tukey HSD test, *p*-value > 0.99). Overall, the bridge strain appears to have destabilized the community via its effect on SBW25, and the plasmids could in some cases exacerbate this effect, resulting in loss of species diversity from the community.

### Plasmids were not always rescued by the corresponding stress

Communities which contained plasmids had increased robustness to environmental stressors to which those plasmids conferred resistance. We therefore expected plasmids to increase in frequency within host populations exposed to a stressor against which the plasmid confers resistance (*i*.*e*., positive plasmid selection (41)). However, we did not consistently observe this. For pQBR57, which encodes resistance to mercury, dynamics were determined by the hosting strain, presence of the bridge strain in the community, and environmental conditions (Figure 3A-D; ANOVA, F_[1,63]_ = 32.41 strain, F_[1,63]_ = 23.32 bridge, F_[3,63]_ = 25.28 stress, F_[3,63]_ = 21.98 bridge × stress interaction, *p*-values < 0.0001). As expected, plasmid frequency declined in the absence of selection, likely owing to fitness cost. While mercury selection was able to prevent decline of pQBR57 (Figure 3B; post-hoc Tukey HSD tests, *p*-values < 0.0001), this was true only for the 2-member community, where SBW25 was the only available host (Figure 3B, circles; post-hoc Tukey HSD tests, *p*-values < 0.0001). The presence of the bridge strain removed any benefit provided by environmental selection (Figure 3B; post-hoc Tukey HSD tests, comparison between SM/p and SMK/p for Hg, *p*-value < 0.0001), and pQBR57 declined similarly across all environmental treatments (Figure 3A-D, triangles; post-hoc Tukey HSD tests, *p*-values > 0.3). Loss of pQBR57 from the community under mercury conditions after day 6 (Figure 3B, triangles) suggests an interplay between bacteria-plasmid and bacteria-bacteria interactions, which we investigated further (next section).

While host strain had a significant impact on pKJK5 (Figure 3E-H; ANOVA, F_[1,60]_ = 285.03, *p*-value < 0.0001), the overall presence of the bridge strain in the community did not (ANOVA, F_[1,60]_ = 0.072, *p*-value = 0.82). Kanamycin selection had a significant effect on pKJK5 (Figure3G&H; post-hoc Tukey HSD tests comparing kanamycin and non-kanamycin environments, *p*-values < 0.0002). Although there was a quantitative difference in pKJK5 dynamics between the kanamycin and dual selection environments in the three-member (but not the two-member) community, thanks to the contribution by the bridge strain (Figure 3G&H, triangles; *p*-values < 0.0002), the overall dynamics were qualitatively similar. In most cases pKJK5 dynamics followed those of MG1655, its predominant host, with little contribution from the bridge strain.

We expected that dual stress conditions (Figure 3D,H,M) would select for KT2440 hosting both plasmids. Indeed, this is the only condition under which we observed any notable presence of co-hosting cells within the community (Figure 3M). However, their increase in frequency was transient, and by day 10 pQBR57 had apparently been lost from the community, while pKJK5 had reached fixation in both of its hosts (Figure 3H). This may once again be due to the interplay of differing plasmid fitness costs for MG1655 and KT2440, different impacts of mercury on the two strains, and competition between them.

### pQBR57 loss under mercury conditions indicates interplay between bacteria-plasmid and bacteria-bacteria interactions

The unexpected loss, only under mercury conditions, of both strains able to carry the plasmid which conferred resistance to mercury (Figure 2B&K) led us to hypothesize that presence of mercury increased the cost of the plasmid (42), exacerbating the competition between SBW25 and KT2440 and driving strain extinction. To test this, we conducted growth curves of SBW25, SBW25 + pQBR57, KT2440, KT2440 + pQBR57, and MG1655 across a range of mercury concentrations (Figure 4A). A comparison of the AUC for the plasmid-free strains shows that increasing mercury concentration has a greater impact on growth of SBW25 and KT2440 than on MG1655, indicating that MG1655 is better able to tolerate mercury stress up to and including 10 µM mercury (Figure 4B), which explains MG1655’s survival under mercury stress. Next, by examining the total frequency of pQBR57 across both hosts (including KT2440 also hosting pKJK5) we found a decline in the non-stressful environment, consistent with pQBR57’s known fitness cost on these strains, and increases in the mercury environment (Figure 4C). Therefore, we propose that in the mercury environment, the toxicity to SBW25 and KT2440 means that only cells harbouring the burdensome plasmid will survive, while they continue to face competition from MG1655, which is able to tolerate the stress without carrying a costly plasmid. This combined effect may also explain the predominance of MG1655 in the 3-member community under dual stress conditions (Figure 2H) over KT2440 (Figure 2M), whose population was ultimately dominated by cells hosting pKJK5 alone (Figure 3H&M). Overall, these results indicate that the combination of host-plasmid-environment interactions and inter-strain competition together shape community outcomes.

**Figure 4.**
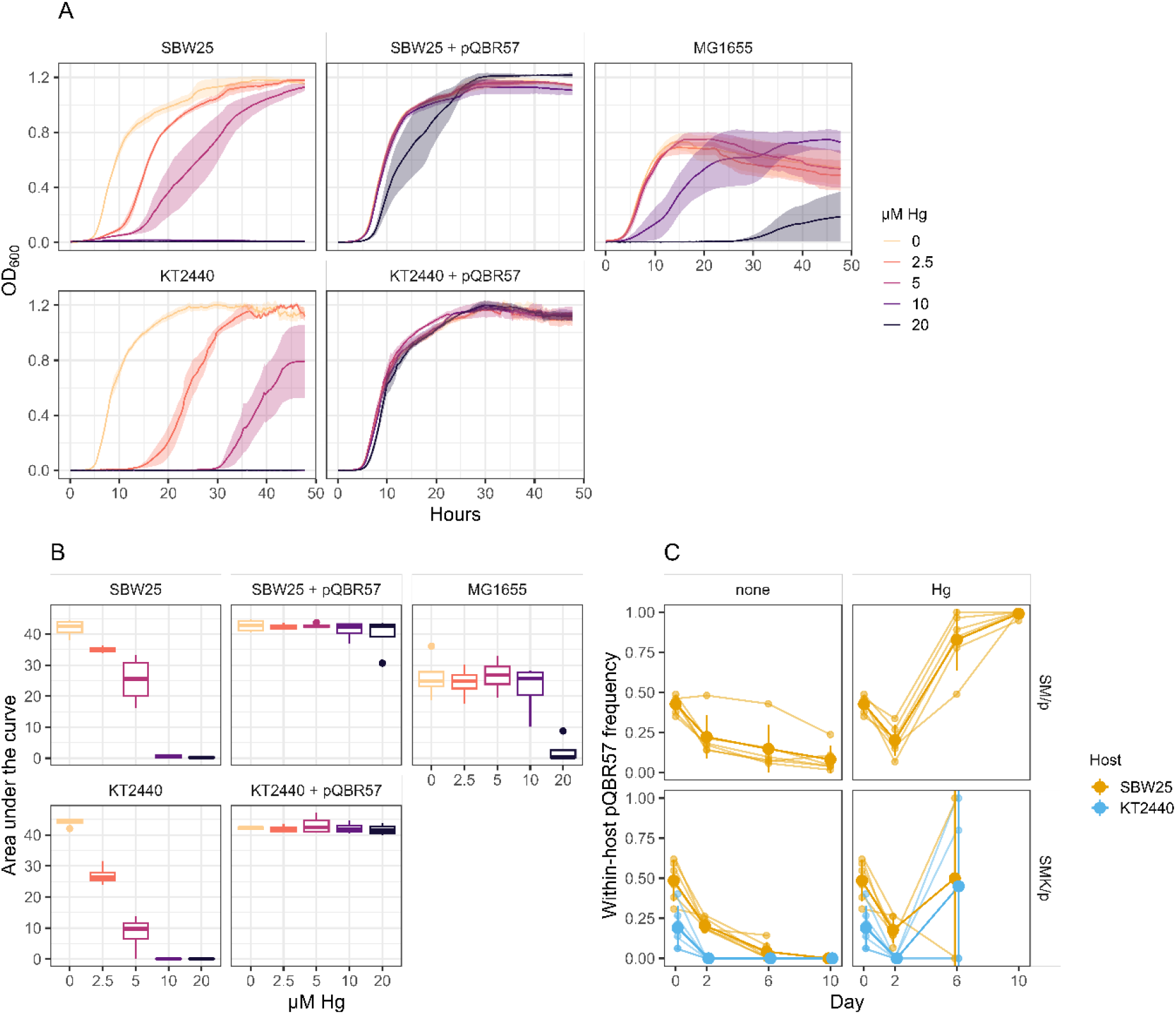
Interplay among strain competition, plasmid hosting, and environment under mercury conditions. We tested the effect of mercury on growth of the three plasmid-free strains and the two pQBR57-hosting strains. (A) Kinetic growth curves of the five strains across a range of mercury concentrations. Solid lines indicate averages of 4 replicate freezer-stock inoculations and dashed lines with shaded areas indicate standard error. (B) Area under the curve of the growth curves in panel A. Boxes show the median (middle line) and the first and third quartiles, whiskers show the smallest and largest value no farther than 1.5 × the inter-quartile range, and dots show outliers). (C) Frequency of pQBR57 within SBW25 and KT2440 under non-stressful and mercury conditions, showing the effect of mercury on plasmid prevalence within each strain. Difference from Figure 3: here KT2440 cells carrying both plasmids are included when quantifying pQBR57 frequency.

## Discussion

While interactions among members shape community diversity, function, and stability, our understanding of the relative roles and cumulative impacts of different types of interactions (*e*.*g*., between bacterial species or between a bacterium and a plasmid) remains limited. This gap hinders our ability to predict the responses of microbial communities to environmental change. Here, we explored the interplay of several forces — inter-species interactions, mobile genetic elements, and environmental change — on microbial community composition and dynamics. We found that interactions between organisms, known to have a predominant role in community dynamics, fundamentally impacted the success of their resident MGEs. Such effects could even overwhelm any benefits provided by the MGEs in relevant stressful environments. For example, we found that the mercury resistance plasmid pQBR57 was lost even from environments containing mercury because its resident hosts were outcompeted, while *E. coli* MG1655 persisted (allowing maintenance of its plasmid pKJK5) regardless of environmental selection pressures. Such patterns challenge our intuition as to drivers of MGEs in communities, indicating that MGE dynamics are fundamentally determined by internal community interactions, and not only by extrinsic selective pressures.

We manipulated community composition by the addition of a bridge strain, *P. putida* KT2440, that could harbour both plasmids. The main effect of the bridge strain on community dynamics was through competition with *P. fluorescens* SBW25, which suffered from reduced abundances across environmental and plasmid conditions (Figure 2A-D). Previous work found that competitive interactions between these species exerted only a mild effect on abundances in a structured soil environment (43) or in heterogenous liquid culture, and though the effects were intensified in well-mixed rich media ((40), as used in the current study), they tended not to result in local extinction of SBW25. Additional competitive pressure from the other community member, MG1655, may have been sufficient to exclude SBW25 from stressful or bridge-strain-containing conditions in our experiments; such competition may be further intensified in minimal media, as opposed to the rich medium used here (44). Natural communities, evolving to specialize on a single labile carbon source, tend to converge on a conserved composition at the Family level, consisting principally of Enterobacteriaceae and Pseudomonadaceae owing to the metabolic capabilities of with each family (44). Similar species-sorting interactions may be at work in our simpler communities, resulting in the exclusion of the less competitive Pseudomonad. As we performed our assays in well-mixed rich media, we encourage future work to investigate community outcomes under resource limitation or in structured, soil-like environments which are expected to better approximate natural conditions.

While we expected that the relative success of the bridge strain would promote the persistence of both plasmids, plasmid maintenance was generally reduced in communities containing the bridge strain. Previous work found that *P. putida* KT2440 was a less favourable host than *P. fluorescens* SBW25 for plasmid pQBR57, even if the fitness effects in each strain are similar ((11,45), Figure S3). The plasmid pKJK5 was more costly to the bridge strain than to its peripheral strain *E. coli* MG1655 (Figure S3) and likewise less likely to be maintained by KT2440. Competitive interactions between organisms, and variance in MGE stability across species and strains (possibly depending on co-infection status of the host (14,15)), thus interact to have consequential effects for maintenance and spread of MGEs that could not be predicted from MGE host ranges alone (46). The plasmid dynamics we report here reflect the net balance of transfer, cost, and loss alongside host competition; future work should additionally quantify specific host-plasmid interactions such as conjugation and segregational loss to further explore drivers of MGEs in communities.

We expected that selection for plasmid-borne resistances would have promoted the corresponding plasmid, and the organisms able to host that plasmid. However, our results suggest that selection for plasmid-borne resistances was insufficient to rescue plasmids, or hosts, in a community context — most strikingly in communities treated with mercury, where both hosts of the mercury resistance plasmid went extinct. The toxicity of mercury to SBW25 and KT2440 (Figure 4) selected for cells harbouring the burdensome plasmid while also allowing ongoing competition from MG1655. The combination of host-plasmid-environment interactions and inter-strain competition may therefore counteract environmental selection and determine plasmid maintenance. Higher concentrations of mercury may have rescued pQBR57 (41), suggesting that local plasmid fate may vary across heterogeneous environments.

Embedding bacterial-plasmid interactions in a community has been shown to impact plasmid fate. Walker-Sünderhauf et al. (47) showed that maintenance of pKJK5 in two different microbes isolated from potting soil (species of *Stenotrophomonas* and *Variovorax*) was disrupted in the context of a five-species community, leading to plasmid loss from these organisms. Kottara et al. (32) similarly showed that plasmid prevalence in a focal species could be reduced by the epidemiological dilution effect of living in a multi-species community, as conjugation events were more likely to result in plasmid transfer to non-focal organisms. Besides shaping microbial community composition, strain competition (*i*.*e*., features of the bacterial interaction matrix ‘A’; Figure 1A) can influence subsequent opportunities for HGT, by driving the dynamics of the vectors of gene transfer, sometimes to extinction. The influence of strain competition on plasmid survival might explain why some plasmids may have evolved to manipulate expression of secreted competitive molecules (48,49).

In a reciprocal manner, microbial community dynamics can be profoundly altered in the context of horizontal gene transfer. HGT can have a positive effect on community diversity and microbiome stability by enabling beneficial resistances to transfer across community members, facilitating their survival (50). The transfer of costly genes can also maintain community stability in a process of ‘dynamic neutrality’, reducing variance in growth rates between species and mitigating competition (51), although this process may be influenced by changes in species interactions. Our work suggests that structure in the HGT network (e.g. presence of a bridge strain, creating partly overlapping host ranges) can have a critical effect modulating the impact of HGT on communities, allowing – or preventing – mobile genes to hitch-hike on competitive community members.

The non-*Pseudomonas* species in our experiment, *E. coli* MG1655, offers a contrast. MG1655 was successful at competition regardless of the presence or absence of the bridge strain and was able to maintain pKJK5 successfully. Moreover, MG1655 was successful at surviving in the face of single stresses even when plasmids were absent. Our concentrations of selective agents were determined based on single-species MIC measurements; growth in a multi-species community, together with the large population size and standing genetic diversity gained during the first growth cycle (from which stresses were absent), likely allowed survival beyond these minimal concentrations. Moreover, mercury resistance is achieved by environmental detoxification (52), a ‘public good’ which could benefit other, nearby community members lacking specific mercury resistance genes. Interestingly, MG1655 could survive dual stresses only in communities with plasmids, even though it couldn’t maintain both plasmids itself, suggesting that resistance to one stress improved the capacity of this strain to cope with a heterologous stress, possibly by enabling sufficient population size for *de novo* mutants to emerge.

Our work demonstrates the importance of considering the interplay of key factors in understanding community dynamics, not just the effects of those factors in isolation. Bacterial community members face effects of biotic and abiotic drivers, such as competition and cooperation, plasmid hosting, and environmental stresses, which can interact in ways that impact their survival differently than when each driver is considered alone. Further, we suggest that plasmids are at risk of being lost from a community due to impacts on their hosts, even when environmental conditions would be expected to select for them. This idea suggests additional complexity that may be at play in plasmid evolution.

## Acknowledgements

This work was supported by a Human Frontiers Science Programme Project Grant to SP, MdeD, JPJH (RGY0064/2022). YJW is supported by the Kreitman School of Advanced Graduate Studies, Ben-Gurion University of the Negev. JPJH is supported by an MRC Career Development Award (MR/W02666X/1). We thank Uli Klümper (TU Dresden) for providing pKJK5::GFP, and Mike Bottery (University of Manchester) for E. coli MG1655.

## Notes

### Competing Interest Statement

The authors have declared no competing interest.

### Summary of Updates

Additional experimental results added (new Figure 4 and Supplement); Figures 2 and 3 revised; Supplemental files updated.

