## Supplemental Information for "Species interactions determine plasmid persistence in a 3-member bacterial community"

Contents:

Supplemental methods

Supplemental results

Table S1. Plasmid-hosting bacterial sub-populations

Figure S1. Total community size across environments

Figure S2. Growth in increasing kanamycin concentrations of MG1655 from plasmid-free communities

Figure S3. Plasmid fitness costs across hosts

Figure S4. Strain survival in experimental communities

Figure S5. Stress tolerance of co-cultured strains

Figure S6. MICs under experimental vs MIC-assay conditions

**Supplemental methods**

*Minimum inhibitory concentration (MIC) assays.* To identify appropriate concentrations of environmental stressors to use in the experiment, we conducted an initial MIC assay whereby all eight bacterial strains were grown overnight in LB broth, then diluted 1:500 and added 50 µl bacteria to 50 µl LB broth with varying concentrations of stressor in a round-bottomed microtiter plate, and incubated on the lab bench (20 ºC) overnight (24 hrs). Growth was scored by eye as a binary indicating whether any turbidity was observed. Similar experiments were also performed using KB broth. To match the experimental conditions, we performed the same assay in LB with a total volume of 200 µl, in a flat-bottomed microtiter plate, at 28 ºC shaking, with observations taken at 24 and 48 hrs. To test the effect of co-culture on community mercury and kanamycin MICs, we performed the initial MIC assay and the MIC assay using experimental conditions with the plasmid-free strains in monoculture, pairwise, and three-way co-culture. After 48 hrs, we estimated surviving population sizes in turbid wells by spot plating onto selective square LB agar plates (250 µg/ml streptomycin incubated at 28 ºC for SBW25, 50 µg/ml X-Gal incubated at 37 ºC to distinguish MG1655 and KT2440).

**Supplemental results**

*MG1655 survives stresses in plasmid-free communities*. We tested two hypotheses for the observed lower impact of mercury and kanamycin stress on MG1655 than on SBW25 or KT2440: (i) that the stresses had different impacts on the communities than on individual strains in monoculture, and (ii) that the effect of the stresses was different under different experimental conditions.

To test whether co-culturing the strains provided a degree of protection from stress compared to monoculture conditions, we performed an additional MIC assay including each strain alone, all paired cultures, and the three-way culture. We observed occasional low levels of growth under the lowest concentrations of stress tested (10 µM mercury, 12.5 µg/ml kanamycin), with survival in kanamycin primarily driven by presence of MG1655 (Figure S5) which we confirmed by spot plating on selective media. This is consistent with the hypothesis that MG1655 can rapidly evolve sufficient kanamycin resistance to survive under our experimental conditions even in the absence of specific resistance genes.

We then compared MIC measurements made under experimental conditions (28 ºC, flat-bottomed microtiter plate, 48 hr growth cycle) with those made with our initial MIC assay conditions (20 ºC, round-bottom microtiter plate, 24 hr growth). For all strains except SBW25

+ pQBR57, we observed higher mercury resistance either after longer incubation (48 hrs) or in KB media (Figure S6A; KB has 20 g/L proteose peptone vs. LB’s 10 g/L casein peptone plus a protein contribution from the 5 g/L yeast extract). This suggests that mercury adsorption by peptides and amino acids in the media is a relevant process in our system (Nies 2003; Ajsuvakova et al. 2020), with decreasing mercury bioavailability within each growth cycle creating a pulse-decay pattern of mercury stress, enabling susceptible strains to grow over a longer growth cycle. For kanamycin stress, we observed almost no difference across assays (Figure S6B).

**Table S1. Plasmid-hosting bacterial sub-populations**

| **Strain** | Plasmid(s) |
|---|---|
| SBW25-Sm$^R$*lacZ* | pQBR57::tdTomato |
| MG1655 | pKJK5::GFP |
| KT2440-Gm$^R$ | pQBR57::tdTomato |
| KT2440-Gm$^R$ | pKJK5::GFP |
| KT2440-Gm$^R$ | pQBR57::tdTomato and pKJK5::GFP |

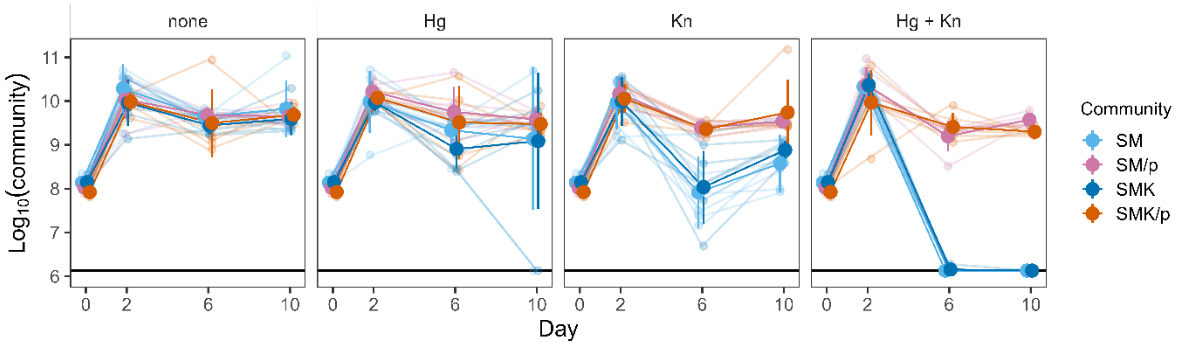


**Figure S1. Total community size across environments.** The total size of all bacterial populations in each community over 10 days, as affected by the environmental conditions. Small dots represent replicate values, large dots represent means of 6 biological replicates, and vertical lines represent 95% confidence intervals of the means.

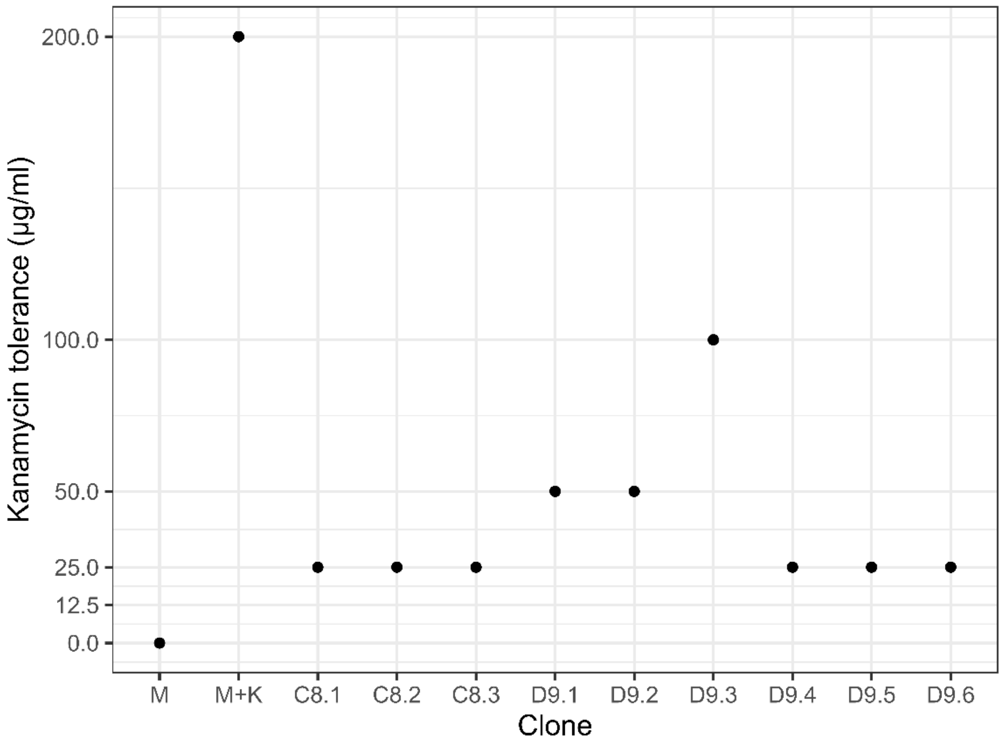


**Figure S2. Growth in increasing kanamycin concentrations of MG1655 from plasmid-free communities.** We isolated clones from dilution plates showing growth of MG1655 under Kn conditions, in communities without plasmids. Dots show the highest kanamycin concentration in which we observed growth. M = MG1655 (negative control), M+K = MG1655 + pKJK5 (positive control).

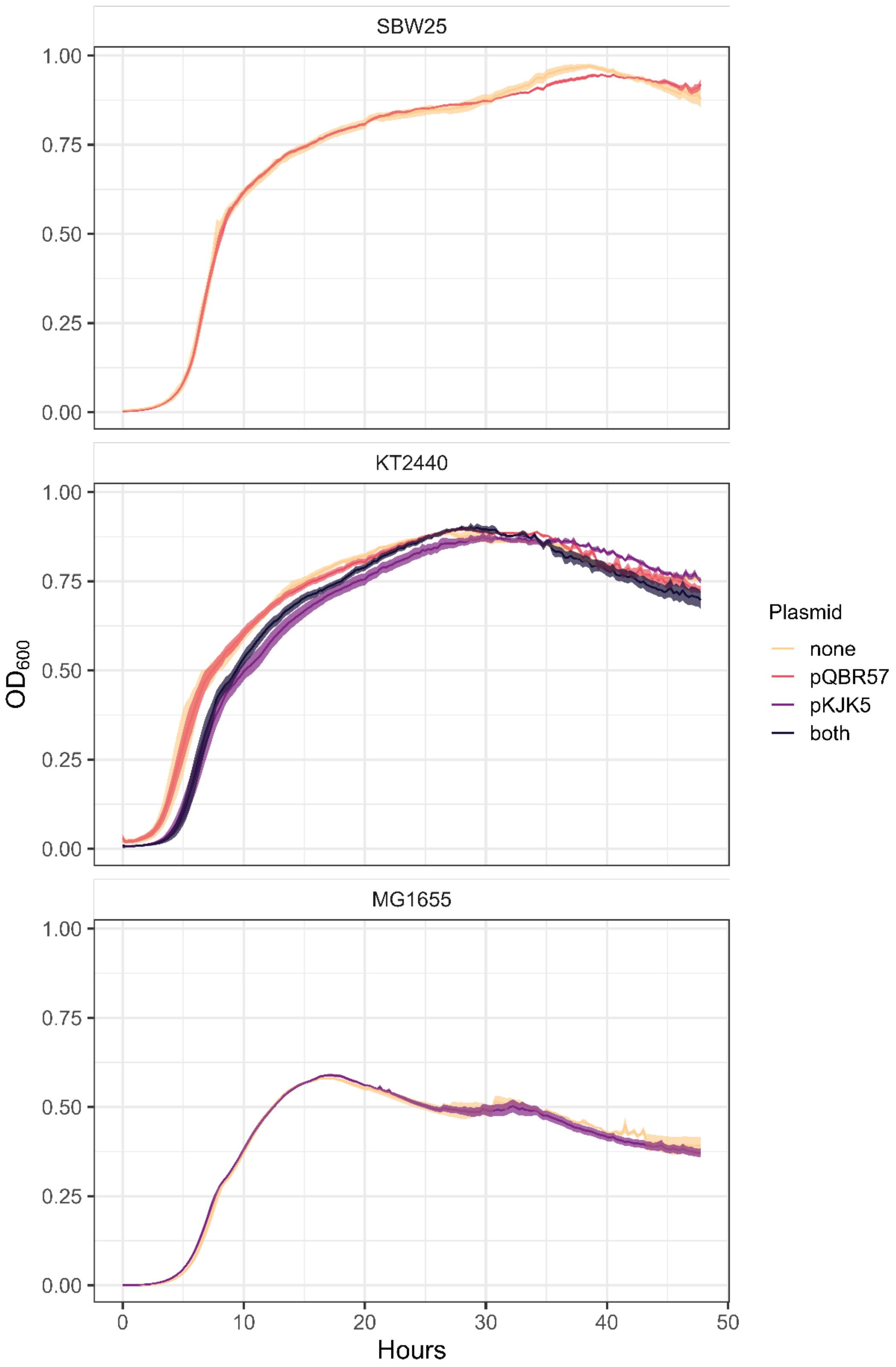


**Figure S3. Plasmid fitness costs across hosts.** Growth profiles of all possible subpopulations (each bacterial strain with each possible plasmid status) over 48 hours, showing the effect of the plasmids on host growth dynamics. Each time step represents 15 minutes. There are 3 biological replicates of each subpopulation.

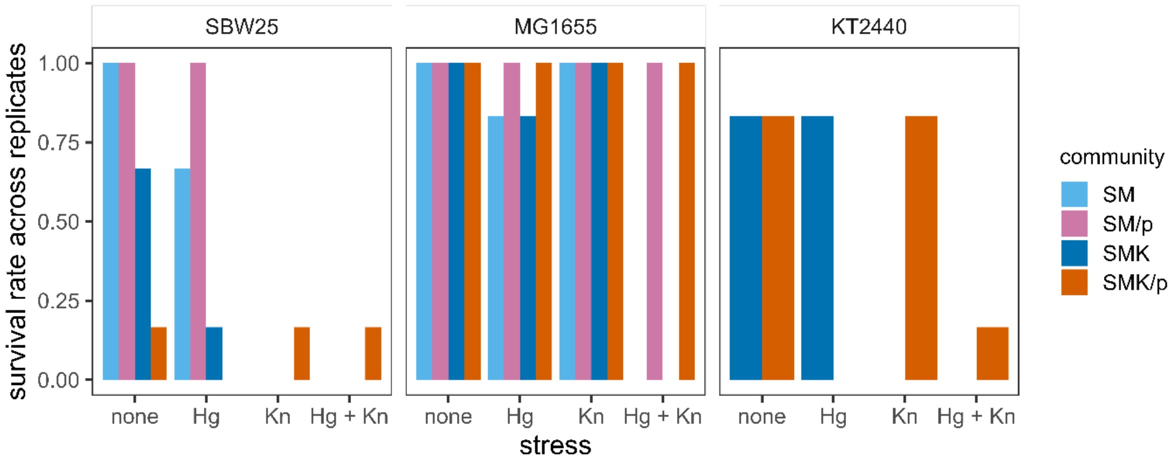


**Figure S4. Strain survival in experimental communities.** Fraction of replicates, out of six total replicates, in which each strain was detected at the final timepoint, across communities and environmental conditions. Limit of detection was 5 x $10^5$ CFUs.

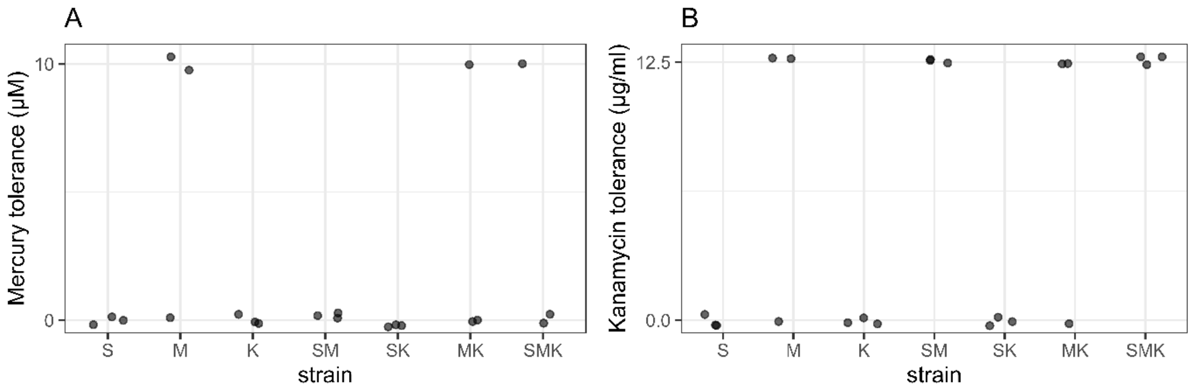


**Figure S5. Stress tolerance of co-cultured strains.** We assayed the ability of plasmid-free strains to grow when co-cultured with each other either pairwise or three-way under experimental conditions. Three replicates are shown; dots are jittered to aid visibility. Mercury was assayed at 10, 20, 40, 80, and 160 μM. Kanamycin was assayed at 12.5, 25, 50, 100, 200 μg/ml. Only the lowest concentrations are shown because no growth was observed in higher concentrations.

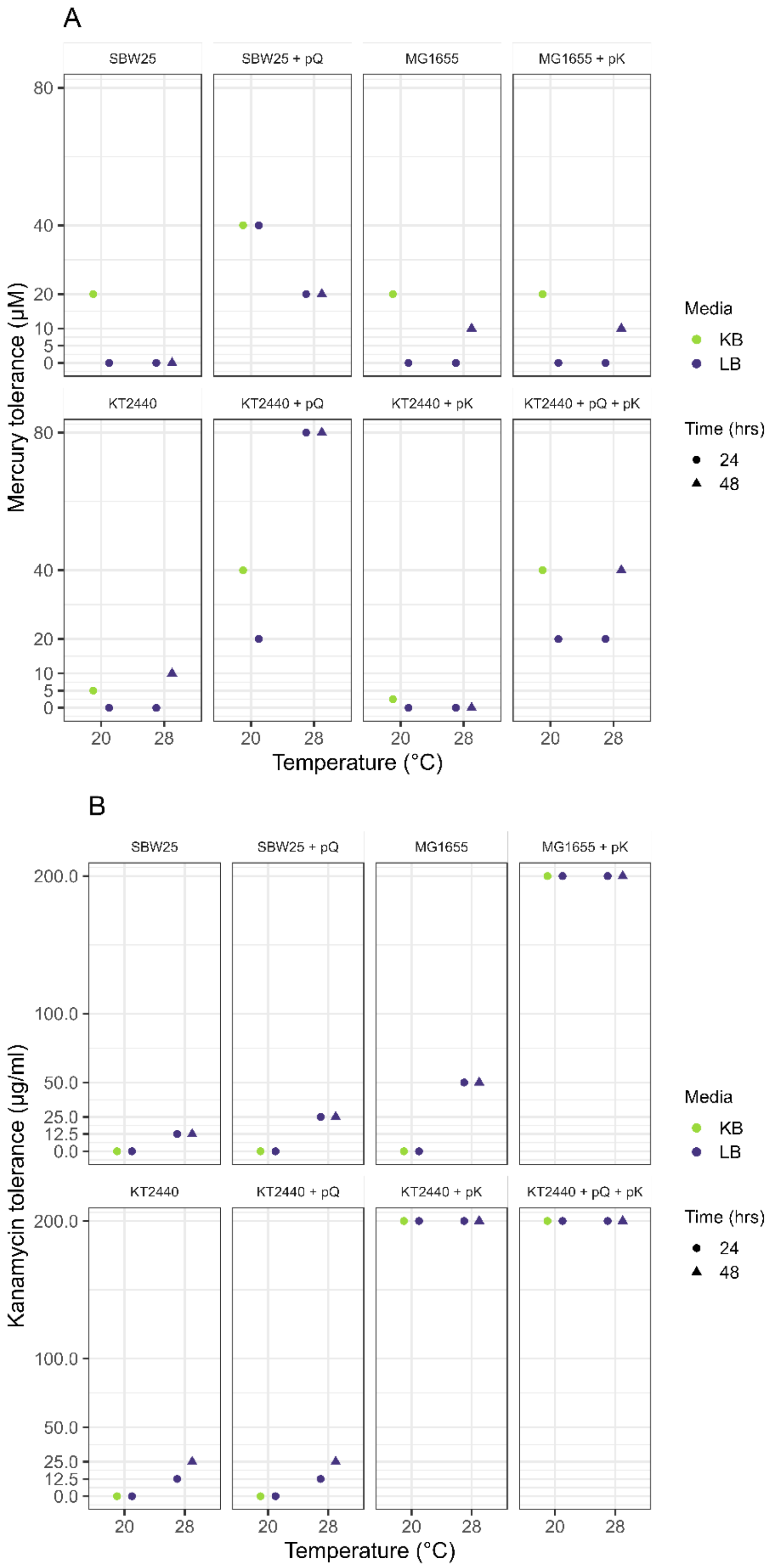


**Figure S6. MICs under experimental vs MIC-assay conditions.** We tested the effect of different assay conditions on the observed mercury and kanamycin MICs of our eight strains. Samples at 20 ºC are from standard MIC assays, and samples at 28 ºC are from assays under experimental conditions (see Methods). Plasmid names are truncated for space: pQ = pQBR57 and pK = pKJK5.
